# The recognition of YopJ family effectors depends on ZAR1/JIM2 immune complex in *Nicotiana benthamiana*

**DOI:** 10.1101/2025.06.04.657961

**Authors:** Injae Kim, Jieun Kim, Ye Jin Ahn, Kee Hoon Sohn, Cécile Segonzac

## Abstract

Pathogens deploy effector proteins to manipulate host physiology and promote infection. YopJ family effectors are highly conserved across bacterial genera that cause crop diseases. Nucleotide-binding leucine-rich repeat receptors (NLRs) play a central role in direct or indirect recognition of effectors and trigger immune responses, including hypersensitive cell death (HR). Two NLRs, *Nicotiana benthamiana* homologs of Pseudomonas tomato race 1 (NbPtr1) and HOPZ-ACTIVATED RESISTANCE 1 (NbZAR1) were recently identified as independently recognizing two YopJ family effectors, HopZ5 and AvrBsT/XopJ2. NbZAR1 also detects XopJ4 via the receptor-like cytoplasmic kinase XOPJ4 IMMUNITY 2 (JIM2). Here, we conducted *Agrobacterium*-mediated transient expression assays with 20 YopJ family effectors from five phytopathogenic bacterial genera and identified 12 YopJ family effectors that are recognized either by NbZAR1 or independently by NbZAR1 and NbPtr1. Furthermore, we show that YopJ family effector-induced HR was differentially suppressed by the deacetylase SUPPRESSOR OF AVRBST-ELICITED RESISTANCE 1, suggesting more than one mechanism for YopJ family effector recognition. This work provides the genetic basis of the recognition of YopJ family effectors in *N. benthamiana* and lays a foundation for the mechanistic study of NbZAR1/JIM2 and NbPtr1 mode of activation.

## Main text

Significant yield losses in a wide range of crop species are due to bacterial diseases (Savary et al., 2019). The type III secretion system (T3SS) is a major virulence determinant of phytopathogenic bacteria. This syringe-like apparatus enables direct translocation of type III effector proteins (T3Es) into host cells (Coll and Valls, 2013). Once delivered, T3Es manipulate host physiology by targeting and modifying specific host proteins (Landry et al., 2020; Peeters et al., 2013; Sabbagh et al., 2019). The Yersinia outer protein J (YopJ) family represents one of the most widely distributed groups of T3Es, present in *Pseudomonas syringae*, *Xanthomonas* spp., *Ralstonia solanacearum*, *Erwinia amylovora*, and *Acidovorax citrulli* (Lewis et al., 2011; Ma and Ma, 2016). YopJ family effectors belong to the C55 protease family, characterized by conserved catalytic residues (Orth et al., 2000). Given the mechanistic similarities between cysteine proteases and acetyltransferases, some YopJ effectors exhibit bifunctional enzymatic activities (Cheong et al., 2014; Ma et al., 2006; Orth et al., 2000). Phytopathogenic bacteria employ YopJ family effectors to enhance virulence in host plants. The *P. syringae* effector HopZ1a disrupts the host microtubule network by acetylating tubulin, inhibiting secretion of antimicrobials (Lee et al., 2012). The *X. euvesicatoria* effector AvrBsT/XopJ2 suppresses AvrBs1-triggered cell death and acetylates ACETYLATED INTERACTING PROTEIN1, a microtubule-associated protein involved in antibacterial immunity (Cheong et al., 2014; Szczesny et al., 2010). Other YopJ family members, such as HopZ4 and XopJ1, interfere with host protein secretion pathways (Üstün et al., 2013; Üstün et al., 2014). Additionally, nuclear-localized YopJ homologs such as XopJ6 and RipP2/PopP2 acetylate WRKY transcription factors to suppress plant defense responses (Lauber et al., 2024; Le Roux et al., 2015; Sarris et al., 2015). Immunosuppressive functions of YopJ family effectors likely underlie the broad conservation of these virulence factors in pathogenic bacteria.

Plants can indirectly detect pathogenic bacteria by monitoring chemical modifications of T3E targets. Modifications of the so-called host guardee/decoy (HGD) activate nucleotide-binding leucine-rich repeat receptors (NLRs) and elicit calcium-dependent immune responses, including reactive oxygen species accumulation, phospho-relay, and transcriptional reprogramming (Kim H. et al., 2023; Ngou et al., 2022). NLR-triggered immunity often culminates in the hypersensitive response (HR), a form of rapid programmed cell death that can limit pathogen spread (Bi et al., 2021; Gao et al., 2013). The *Arabidopsis* coiled-coil NLR (CNL) HOPZ-ACTIVATED RESISTANCE 1 (ZAR1) is one of the best-characterized plant NLRs and participates in the indirect recognition of diverse T3Es (Laflamme et al., 2020; Lewis et al., 2010). ZAR1 monitors modifications of receptor-like cytoplasmic kinases (RLCKs) to detect effector activity (Ma et al., 2006; Martel et al., 2020; Seto et al., 2017; Wang et al., 2019).

In *Arabidopsis*, the *P. syringae* effector HopZ1a directly acetylates the RLCK-XII pseudokinase HopZ-ETI-Deficient1, which interacts with ZAR1 and triggers immune activation (Lewis et al., 2013). Similarly, the *Nicotiana benthamiana* homolog of ZAR1 (NbZAR1) recognizes the *X. perforans* effector XopJ4 via the RLCK-XII protein XOPJ4 IMMUNITY 2 (JIM2) (Schultink et al., 2019). NbZAR1 also detects the *X. euvesicatoria* effector AvrBsT/XopJ2 and the *P. syringae* pv. *actinidiae* effector HopZ5 (Ahn et al., 2023). Both HopZ5 and AvrBsT/XopJ2 are also independently recognized by NbPtr1, the *N. benthamiana* homolog of Pseudomonas tomato race 1 resistance protein, revealing that individual T3E can be perceived by non-orthologous NLRs within plant species (Ahn et al., 2023; Kim H. et al., 2023; Mazo-Molina et al., 2020).

Previous studies reported that other YopJ family effectors can induce cell death in *N. benthamiana*. (Figure S1) (Albers et al., 2019; Ma et al., 2006; Thieme et al., 2007; Traore et al., 2019; Vinatzer et al., 2006). Here, we used *Agrobacterium* to transiently express 20 representative YopJ family T3Es in *N. benthamiana* plants that are knocked down or knocked out for *NbZAR1* and/or *NbPtr1* to demonstrate the genetic basis of YopJ family T3E recognition. We found that 12 YopJ family T3Es are indirectly recognized through their predicted enzymatic activity. Additionally, we report differential suppression of NbZAR1/JIM2- and NbPtr1-mediated HR by the deacetylase SUPPRESSOR OF AVRBST ELICITED RESISTANCE 1 (SOBER1) (Bürger et al., 2017; Choi et al., 2021; Cunnac et al., 2007). The specificity of YopJ family T3E recognition by NbZAR1 and/or NbPtr1 aligns with phylogenetic grouping, providing mechanistic insights for the mode of activation of two immune receptors, which could aid in crop protection against diverse bacterial diseases.

We selected 20 YopJ family T3Es based on annotations in the T3E repertoires of *P. syringae*, *Xanthomonas* spp., *R. solanacearum*, *A. citrulli*, and *E. amylovora* (Figure S1) (Ahn et al., 2023; Lauber et al., 2024; Ma and Ma, 2016; Peeters et al., 2013). The corresponding sequences harbor predicted catalytic residues characteristic of the YopJ family and cofactor inositol hexaphosphate (InsP6)-binding site, except RipJ and HopZ1c, respectively (Figure 1a) (Ma et al., 2015; Pandey et al., 2021; Xia et al., 2021; Zhang et al., 2016; Zhou et al., 2009). The RLCK-XII family protein, JIM2, is essential for NbZAR1 activation and is expressed at a low level in *N. benthamiana* (Figure S2) (Ahn et al., 2023; Kurotani et al., 2025; Schultink et al., 2019). For this reason, we monitored the cell death induced by YopJ family T3Es co-expressed with JIM2 or GFP. We used XopAU, an unrelated *X. euvesicatoria* T3E, as a positive control for cell death (Figure 1b) (Teper et al., 2018). Cell death intensity was quantified by measuring chlorophyll fluorescence parameters (quantum yield, QY) in the agro-infiltrated tissues (Figure S3) (Kim B. et al., 2023). In Nb-1 TRV:EV plants, at 3 days post-infiltration, we observed three categories of responses. Aave2166, AvrBsT/XopJ2, and HopZ5 form a first group (group I) that triggered a robust cell death when co-expressed with GFP, regardless of additional expression of JIM2. A second group (group II) comprising HopZ1a, HopZ1b, HopZ2, HopZ4, XopJ1, XopJ3, XopJ4, RipP1, and Aave2708 induced cell death only when co-expressed with JIM2. Lastly, HopZ1c, HopZ3, XopJ5, XopJ6, RipJ, RipK, RipP2, and Eop1 caused weak or no cell death regardless of JIM2 expression and were categorized into group III (Figures 1b and S3). As we confirmed the accumulation of all the YopJ family T3E proteins by immunoblotting (Figure S4), the lack of response in tissues expressing group III T3Es can be interpreted as a lack of recognition in *N. benthamiana*.

**Figure 1.**
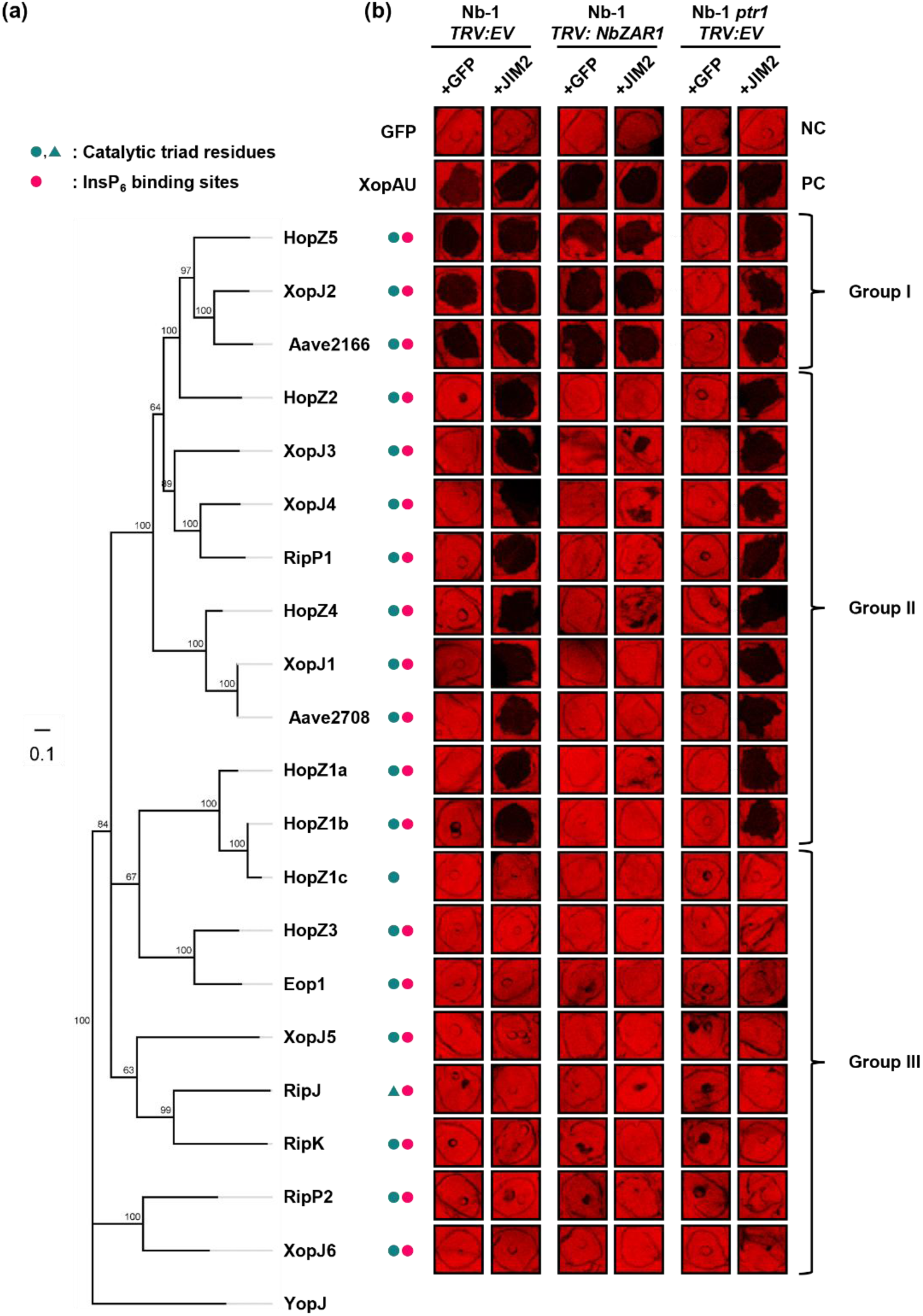
Twelve YopJ family effectors trigger NbPtr1- and/or NbZAR1/JIM2-dependent cell death in *Nicotiana benthamiana*. (a) Phylogenetic tree of 21 YopJ family effectors. The consensus tree was constructed based on multiple sequence alignment using Clustal Omega 1.2.2 with bootstrap resampling. Genetic distances were calculated using the Jukes-Cantor model, and the tree was built using the Neighbor-Joining method. *Yersinia* YopJ was used as an outgroup from an animal pathogen. Green circles and triangles indicate conserved catalytic residues (cysteine-based and serine-based, respectively). Pink circles denote the presence of InsP₆ binding sites (Zhang et al., 2016). (b) Green fluorescent protein (GFP), XopAU and each YopJ family effector C-terminally fused with yellow fluorescent protein (YFP) (OD_600_ = 0.5) were co-expressed with P19 (OD_600_ = 0.1) and either GFP (OD_600_ = 0.5) or JIM2-4xMyc (OD_600_ = 0.5) in *N. benthamiana* wild-type plants (Nb-1) silenced for empty vector (*TRV: EV)* or *NbZAR1* (*TRV: NbZAR1)* or in *ptr1* knock-out plants (Nb-1 *ptr1 TRV: EV*). NC indicates a negative control, and PC indicates a positive control of the experiment. Infiltrations were photographed in light-emitting diode (LED) light (red-orange 617 nm and cool white 6500 K) at three days post-infiltration (dpi). The cell death was quantified by measuring chlorophyll quantum yield (QY) (Figure S3). This experiment was independently repeated twice with at least three replicated spots. Group I effectors are recognized by both NbPtr1 and NbZAR1. Group II effectors are recognized by NbZAR1 only. Group III effectors do not trigger cell death.

In the presence of JIM2, the cell death induced by group II T3Es was lost in Nb-1 plants silenced for *NbZAR1* (Figures 1b and S3). Hence, our analysis demonstrates that 12 out of the selected 20 YopJ family effectors, including T3Es that are previously reported to trigger cell death in *N. benthamiana*, are recognized by NbZAR1/JIM2 (Ahn et al., 2023; An et al., 2025; Ma et al., 2006; Schultink et al., 2019). Furthermore, since XopJ2 and HopZ5 (group I) are independently recognized by NbPtr1 (Ahn et al., 2023), we expressed the 20 YopJ family T3Es in Nb-1 *ptr1*, a *NbPtr1* knock-out line generated using CRISPR/Cas9 technology (Ahn et al., 2023). HopZ5, XopJ2, and Aave2166 (Group I) co-expressed with GFP could not induce cell death in Nb-1 *ptr1*, but co-expression with JIM2 restored intense cell death (Figures 1b and S3). Additionally, we confirmed that Group I YopJ T3Es failed to induce cell death in Nb-1 *ptr1* TRV: *NbZAR1* plants (Figures 1b and S5). Thus, Aave2166 from *A. citrulli*, AvrBsT/XopJ2 from *X. euvesicatoria,* and HopZ5 from *P. syringae* are independently recognized by NbZAR1 and NbPtr1. Notably, the YopJ family effectors recognized by NbZAR1 and/or NbPtr1 are phylogenetically close to one another, and group I effectors form a subclade, indicating that YopJ effector recognition in *N. benthamiana* depends on conserved activity or HGDs that the effectors target.

NLRs can sense the presence of effectors through monitoring effector activities in a host cell (Jones et al., 2024). The 20 selected YopJ effectors harbor a conserved catalytic triad including cysteine, histidine, and either aspartate or glutamate residues, with the exception of RipJ (Figure 1a) (Pandey et al., 2021). To assess the role of the T3E enzymatic activity for the activation of NbZAR1/JIM2 and NbPtr1, we generated cysteine-to-alanine substitution (C/A) mutants for the three effectors from group I and the nine effectors from group II that induce NLR-dependent cell death. This substitution of the conserved cysteine was shown to impair the catalytic activity and the recognition of HopZ1a and RipP2 (Lee et al., 2012; Tasset et al., 2010). The 12 C/A mutants induced significantly reduced or no cell death compared to their wild-type counterparts in Nb-1 plants (Figures 2 and S6). Since all C/A mutants accumulated to detectable protein levels (Figure S7), these results indicate that the predicted catalytic activity of the group I and group II effectors is required for the recognition by NbZAR1 and/or NbPtr1, consistent with the previously established indirect mode of activation of these NLRs (Ahn et al., 2023; Mazo-Molina et al., 2020; Schultink et al., 2019).

**Figure 2.**
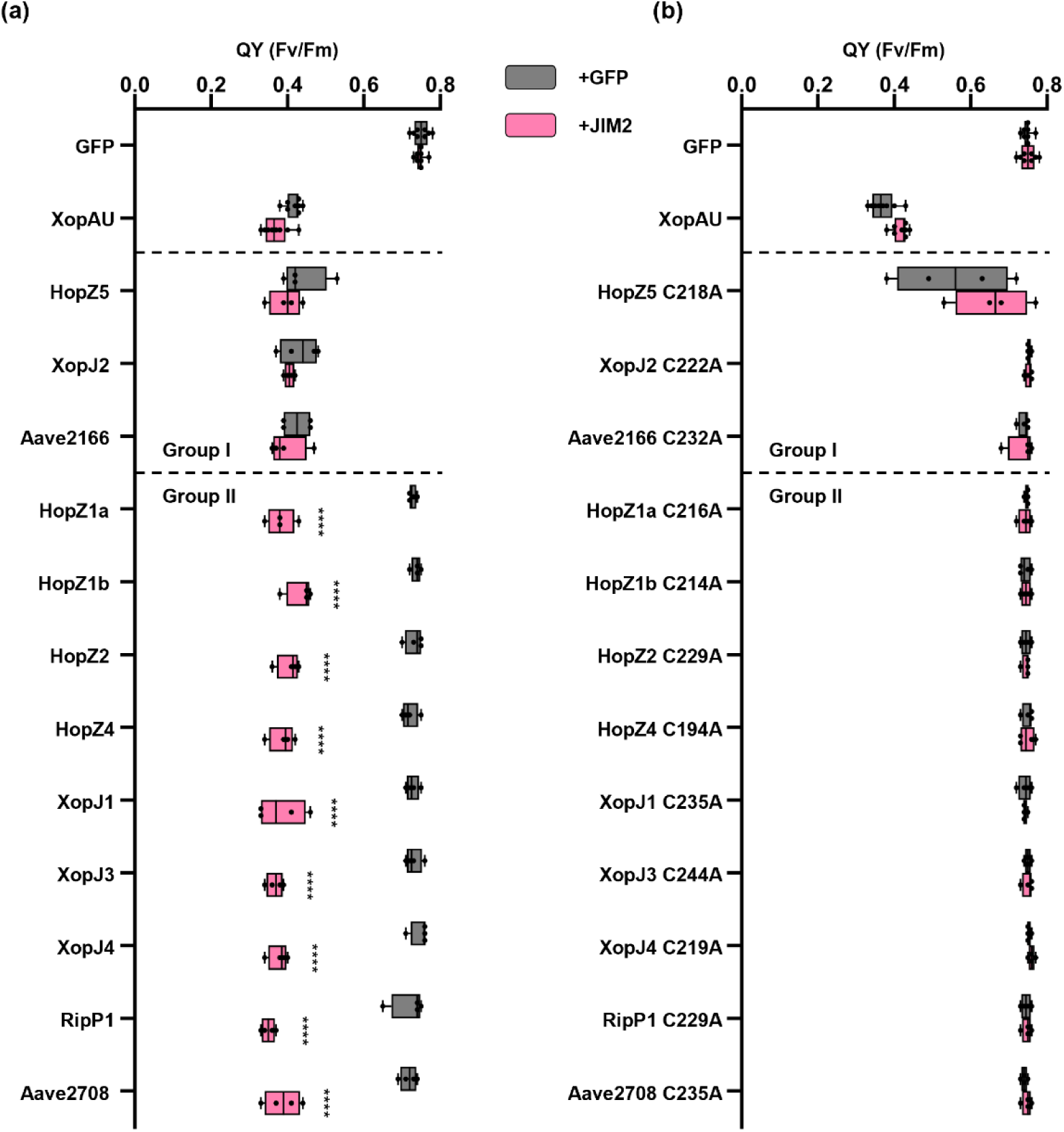
The conserved catalytic cysteine of YopJ family effectors is required for recognition by NbPtr1 and NbZAR1. GFP, XopAU, and YopJ family effectors (a) or their corresponding catalytic cysteine mutants (C/A) (b) were co-expressed with P19 (OD_600_ = 0.1) and with GFP (OD_600_ = 0.5) or JIM2-4xMyc (OD_600_ = 0.5) in *N. benthamiana* wild-type plants (Nb-1). Infiltrated leaves were photographed in light-emitting diode (LED) light at 3 dpi (Figure S6). Cell death intensity was quantified by measuring the quantum yield (QY) of each infiltrated spot. High QY indicates strong cell death at the infiltration site, and low QY indicates weak or no cell death. Box plots show the distribution of individual values between the lower and upper quartiles (25-75%), individual values (dots), and median value (line). The whiskers indicate minimum and maximum values. Statistical analysis was performed using two-way ANOVA, with JIM2 co-expression as the main factor and effector identity included as a blocking factor to account for grouped comparisons. Dunnett’s multiple comparison test was applied post hoc. Asterisks indicate statistically significant differences between the co-expression of GFP and JIM2 (****, *P* < 0.0001). This experiment was independently repeated twice.

While a protease activity has been reported at low levels for some YopJ family effectors (Ma et al., 2006), subsequent studies reported that YopJ family effectors modify their host targets via acetylation (Cheong et al., 2014; Choi et al., 2021; Jeleńska et al., 2021; Le Roux et al., 2015; Lee et al., 2012; Rufián et al., 2021). The *Arabidopsis* deacetylase SOBER1 was shown to deacetylate the substrate of HopZ5 and consequently abolish HopZ5-induced cell death (Burger et al., 2017; Choi et al., 2021). To further assess the relationship between the acetyltransferase activity of YopJ family effectors and their recognition by NbPtr1 and NbZAR1, we co-expressed the 12 recognized YopJ family effectors with SOBER1 in Nb-1 and Nb-1 *ptr1* plants (Figures 3 and S8). First, we observed differential responses in the presence of SOBER1 and re-categorized group II T3Es into two sub-groups. In Nb-1 plants, in the presence of JIM2, the cell death induced by HopZ4, XopJ1, and Aave2708 (group IIa) was completely suppressed by SOBER1, while the cell death induced by HopZ1a, HopZ1b, HopZ2, XopJ3, XopJ4, and RipP1 (group IIb) was not. Interestingly, these findings suggest that group IIa and group IIb T3Es may modify multiple host proteins, including JIM2, while only some of these could be substrates for SOBER1 deacetylase activity. The substrate specificity of SOBER1 has been previously reported in *N. tabacum*, where SOBER1 can suppress XopJ2/AvrBsT- and HopZ5- but not HopZ1a-triggered cell death (Cheong et al., 2014; Choi et al., 2018; Choi et al., 2021; Jayaraman et al., 2017). Alternatively, group IIa effectors might act as proteases rather than acetyltransferases (Üstün et al., 2014; Üstün and Börnke, 2015), so it remains possible that NbZAR1 monitors a protease activity of group IIa effectors. Given that the substrate preference of SOBER1 can be altered by mutation on the hydrophobic anchoring residues, and that other α/β hydrolases closely related to SOBER1 have different substrate specificities (Bürger et al., 2017), there is a possibility that SOBER1 could suppress a distinct enzymatic activity of these effectors, such as the previously reported protease activity (Üstün et al., 2014; Üstün and Börnke, 2015). It would be worthwhile to investigate whether SOBER1 variants and hydrophobic anchor mutants retain the ability to suppress NbZAR1-mediated recognition.

**Figure 3.**
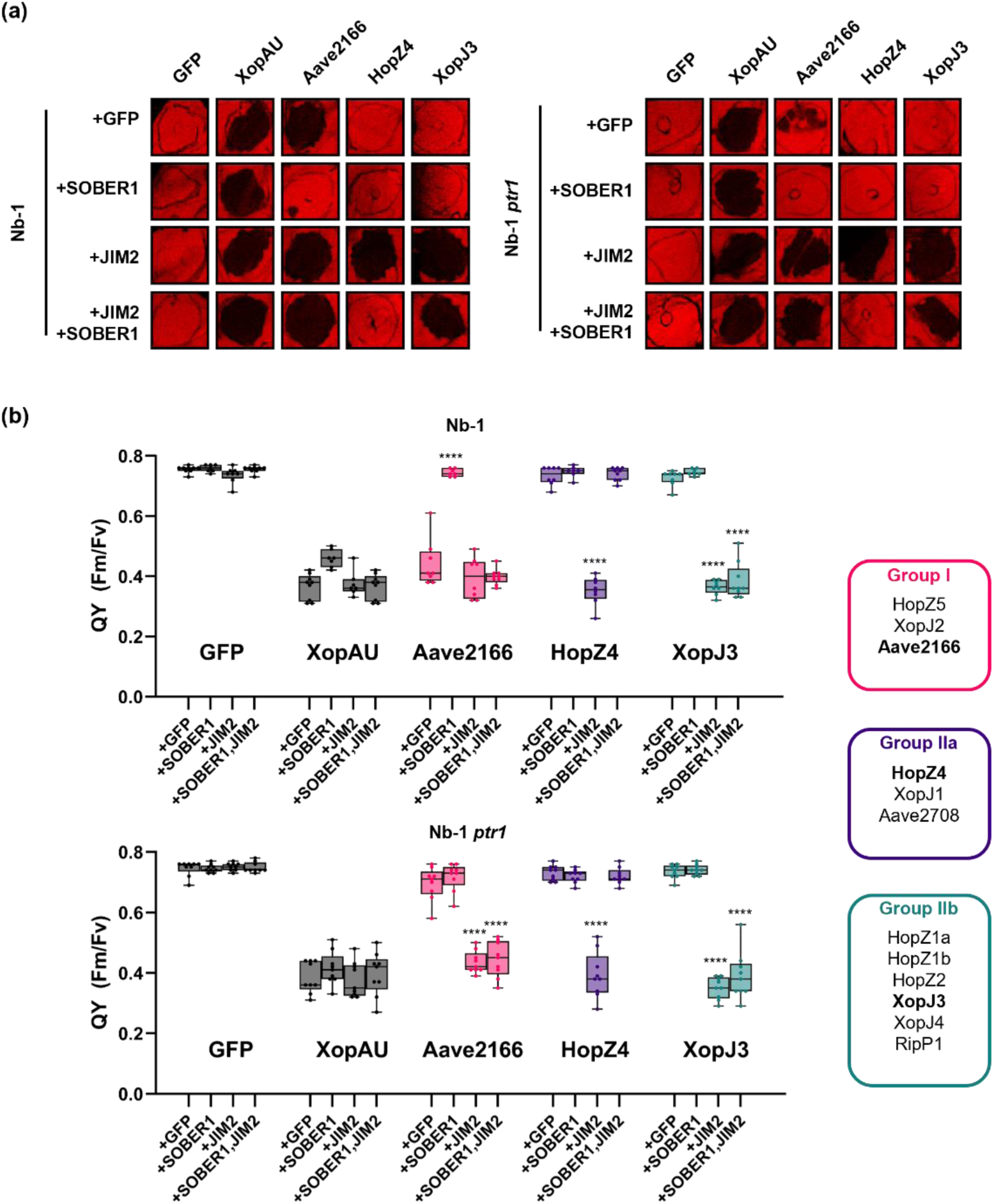
Both NbPtr1- and NbZAR1-dependent recognition of YopJ family effectors can be suppressed by the deacetylase SOBER1. (a) GFP, XopAU, Aave2166 (representative of group I), HopZ4 (representative of group II a) and XopJ3 (representative of group IIb) (OD_600_ = 0.1) were co-expressed with P19 (OD_600_ = 0.1) and with GFP (OD_600_ = 0.4), with SOBER1-Flag (OD_600_ = 0.4) and GFP (OD_600_ = 0.1), with JIM2-4xMyc (OD_600_ = 0.1) and GFP (OD_600_ = 0.4), or with JIM2-4xMyc (OD_600_ = 0.1) and SOBER1-Flag (OD_600_ = 0.4) in Nb-1 and Nb-1 *ptr1* plants. Infiltrated leaves were photographed in light-emitting diode (LED) light at 3 dpi. (b) The cell death was quantified by measuring quantum yield (QY) at 3 dpi. High QY indicates strong cell death at the infiltration site, and low QY indicates weak or no cell death. Box plots show the distribution of individual values between the lower and upper quartiles (25–75%), individual values (dots), and the median (line). The whiskers indicate the minimum and maximum values. Statistical analysis was performed using two-way ANOVA, with the co-expressed protein as the main factor and effector identity included as a blocking factor to account for grouped comparisons. Dunnett’s multiple comparison test was applied post hoc. Asterisks indicate statistically significant differences compared to GFP co-expression for each effector (****, *P* < 0.0001). These experiments were independently repeated twice, with at least four replicated spots per condition. Boxes indicate the phenotypic effector groups; data for each of the 12 T3Es are presented in Figure S8.

Unexpectedly, the NbPtr1-dependent cell death triggered by HopZ5, XopJ2, and Aave2166 observed in Nb-1 plants in the presence of GFP was significantly suppressed by SOBER1 (Figures 3 and S8). However, SOBER1 did not suppress the NbZAR1-dependent recognition of these three group I effectors, as observed in Nb-*ptr1* plants co-expressing JIM2. Since the effector protein accumulation was unaffected by the presence of SOBER1 (Figure S9), these results suggest that NbPtr1 could monitor the acetylation of yet unidentified HGD. Additionally, we cannot exclude that in our experimental conditions, SOBER1 expression could have been insufficient to remove all acetyl groups from multiple T3E targets and prevent NbZAR1 activation.

In summary, our cell death assays demonstrate that NbZAR1 can recognize 12 YopJ family T3Es in the presence of the key immune component JIM2, providing a genetic basis for the broad recognition spectrum of NbZAR1 and supporting convergently evolved effector recognition by NbZAR1 and NbPtr1 (Ahn et al., 2023; Kim H. et al., 2023). The differential suppression by SOBER1 across the phylogeny of YopJ family effectors further highlights the distinct recognition modes of NbPtr1 and NbZAR1, possibly mediated by specific targets acting as HGD.

## Supporting information

Kim I et al_supp info

## Author contributions

Conceptualization: KHS, CS; Investigation: IK, YJA, JK; Resources: CS, KHS; Supervision: KHS, CS; Funding acquisition: CS; Writing: IK, CS with inputs from all the authors.

## Acknowledgements

This work was supported by the National Research Foundation of Korea (NRF) funded by the Korean Ministry of Sciences and ICT (Projects No RS-2018-NR031006 and No RS-2024-00349151) and by the Korea Institute of Planning and Evaluation for Technology in Food, Agriculture, and Forestry (IPET) through the Agriculture and Food Convergence Technologies Program for Research Manpower Development, funded by the Ministry of Agriculture, Food and Rural Affairs (MAFRA) (No. RS-2024-00398300). The authors declare no conflict of interest.

## Data availability statement

The data that support the findings of this study are available from the corresponding author upon reasonable request.

## Supporting information legends

Figure S1. Features of the 20 YopJ family effectors selected for this study.

Figure S2. Gene expression profile of *JIM2*, *NbZAR1,* and *NbPtr1*. Expression of *JIM2*, *NbZAR1,* and *NbPtr1* in transcript per million (TPM) was retrieved on the NbenBase gene expression tool (https://nbenthamiana.jp/nbrowser/profile) (Kurotani et al., 2025). Red arrows indicate leaves used for the agro-infiltration experiments.

Figure S3. Quantification of cell death in YopJ family T3E expressing tissues shown in Figure 1a. Cell death intensity was quantified by measuring the quantum yield (QY) of each infiltrated spot. High QY indicates strong cell death at the infiltration site, and low QY indicates weak or no cell death. Box plots show the distribution of individual values between the lower and upper quartiles (25-75%), individual values (dots), and median value (line). The whiskers indicate minimum and maximum values. Statistical comparisons were performed using two-way ANOVA with JIM2 co-expression as a main factor and effector identity included as a blocking factor to account for grouped comparisons, followed by Dunnett’s multiple comparison test. Asterisks indicate statistically significant differences compared with the GFP control co-expressed with GFP or JIM2 for each effector in each type of plant (**, *P* < 0.01; ***, *P* < 0.001; ****, *P* < 0.0001). This experiment was independently repeated twice with at least four replicated spots.

Figure S4. Accumulation of YopJ family T3E proteins in tissues shown in Figure 1.

GFP and effector-YFP fusions were expressed with JIM2-4xMyc in Nb-1 *ptr1 TRV: NbZAR1* tissues. Leaf samples were harvested 40 to 60 hours after agroinfiltration. Immunodetection on total protein extracts was performed with anti-GFP antibodies. Ponceau S staining (PS) shows equal loading of the samples.

Figure S5. HopZ5, XopJ2, and Aave2166 are independently recognized by NbPtr1 and NbZAR1.

(a) GFP, XopAU, and each YopJ family effector were co-expressed with P19 and either GFP or JIM2-4xMyc in Nb-1 *TRV: EV* or in Nb-1 *ptr1 TRV: NbZAR1* silenced plants.

(b) Cell death intensity was quantified by measuring the quantum yield (QY) of each infiltrated spot. High QY indicates strong cell death at the infiltration site, and low QY indicates weak or no cell death. Box plots show the distribution of individual values between the lower and upper quartiles (25-75%), individual values (dots), and median value (line). The whiskers indicate minimum and maximum values. Statistical comparisons were performed using two-way ANOVA with JIM2 co-expression and types of plant as main factors and effector identity included as a blocking factor to account for grouped comparisons, followed by Dunnett’s multiple comparison test. Asterisks indicate statistically significant differences between Nb-1 *TRV: EV* and Nb-1 *ptr1 TRV: NbZAR1* for each effector (****, *P* < 0.0001). This experiment was independently repeated twice with at least 4 replicated spots.

Figure S6. Photographs of infiltrated leaf spots shown in Figure 2.

Leaves were photographed in light-emitting diode (LED) light (red-orange 617 nm and cool white 6500 K) at 3 dpi.

Figure S7. Accumulation of YopJ family T3E C/A mutant proteins in tissues shown in Figure 2. GFP and C/A mutant-YFP fusions were expressed with JIM2-4xMyc in Nb-1 *ptr1 TRV: NbZAR1* tissues. Leaf samples were harvested 40 to 60 hours after agroinfiltration. Immunodetection was performed with anti-GFP and anti-MYC antibodies. Ponceau S staining (PS) shows equal loading of the samples.

Figure S8. Photographs of infiltrated leaf spots shown in Figure 3.

Leaves were photographed in light-emitting diode (LED) light (red-orange 617 nm and cool white 6500 K) at 3 dpi.

Figure S9. Accumulation of YopJ family T3E, JIM2 and SOBER1 proteins in tissues shown in Figure 3. GFP and effector-YFP fusions were expressed with JIM2-4xMyc and SOBER1-3xFLAG in Nb-1 *ptr1 TRV: NbZAR1* tissues. Leaf samples were harvested 40 to 60 hours after agroinfiltration. Immunodetection was performed with anti-GFP, anti-MYC, and anti-FLAG antibodies. Ponceau S staining (PS) shows equal loading of the samples.

## References

Ahn, Y.J., Kim, H., Choi, S., Mazo-Molina, C., Prokchorchik, M., Zhang, N., et al. (2023) Ptr1 and ZAR1 immune receptors confer overlapping and distinct bacterial pathogen effector specificities. New Phytologist, 239, 1935–1953.

An, Y., Lu, J., Zhang, S., Fang, B. & Zhang, M. (2025) New insights into CNL-mediated immunity through recognition of Ralstonia solanacearum RipP1 by NbZAR1. Journal of Integrative Plant Biology, 67, 1220–1222.

Albers, P., Üstün, S., Witzel, K., Kraner, M. & Börnke, F. (2019) A remorin from *Nicotiana benthamiana* interacts with the *Pseudomonas* Type-III effector protein HopZ1a and is phosphorylated by the immune-related kinase PBS1. Molecular Plant-Microbe Interactions, 32, 1229–1242.

Bi, G., Su, M., Li, N., Liang, Y., Dang, S., Xu, J., et al. (2021) The ZAR1 resistosome is a calcium-permeable channel triggering plant immune signaling. Cell, 184, 3528–3541.e12.

Bürger, M., Willige, B.C. & Chory, J. (2017) A hydrophobic anchor mechanism defines a deacetylase family that suppresses host response against YopJ effectors. Nature Communications, 8, 2201.

Cheong, M.S., Kirik, A., Kim, J.-G., Frame, K., Kirik, V. & Mudgett, M.B. (2014) AvrBsT acetylates *Arabidopsis* ACIP1, a protein that associates with microtubules and ss required for immunity. PLOS Pathogens, 10, e1003952.

Choi, S., Jayaraman, J. & Sohn, K.H. (2018) *Arabidopsis thaliana* SOBER1 (SUPPRESSOR OF AVRBST-ELICITED RESISTANCE 1) suppresses plant immunity triggered by multiple bacterial acetyltransferase effectors. New Phytologist, 219, 324–335.

Choi, S., Prokchorchik, M., Lee, H., Gupta, R., Lee, Y., Chung, E.-H., et al. (2021) Direct acetylation of a conserved threonine of RIN4 by the bacterial effector HopZ5 or AvrBsT activates RPM1-dependent immunity in *Arabidopsis*. Molecular Plant, 14, 1951–1960.

Coll, N.S. & Valls, M. (2013) Current knowledge on the *Ralstonia solanacearum* type III secretion system. Microbial biotechnology, 6, 614–620.

Cunnac, S., Wilson, A., Nuwer, J., Kirik, A., Baranage, G. & Mudgett, M.B. (2007) A Conserved carboxylesterase is a SUPPRESSOR OF AVRBST-ELICITED RESISTANCE in *Arabidopsis*. The Plant Cell, 19, 688–705.

Gao, X., Chen, X., Lin, W., Chen, S., Lu, D., Niu, Y., et al. (2013) Bifurcation of *Arabidopsis* NLR immune signaling via ca2+-dependent protein kinases. PLoS Pathogens, 9, e1003127.

Jayaraman, J., Choi, S., Prokchorchik, M., Choi, D.S., Spiandore, A., Rikkerink, E.H., et al. (2017) A bacterial acetyltransferase triggers immunity in *Arabidopsis thaliana* independent of hypersensitive response. Scientific Reports, 7, 3557.

Jeleńska, J., Lee, J., Manning, A.J., Wolfgeher, D.J., Ahn, Y., Walters-Marrah, G., et al. (2021) *Pseudomonas syringae* effector HopZ3 suppresses the bacterial AvrPto1–tomato PTO immune complex via acetylation. PLOS Pathogens, 17, e1010017.

Jones, J.D.G., Staskawicz, B.J. & Dangl, J.L. (2024) The plant immune system: From discovery to deployment. Cell, 187, 2095–2116.

Kim, B., Kim, I., Yu, W., Li, M., Kim, H., Ahn, Y.J., et al. (2023) The *Ralstonia pseudosolanacearum* effector RipE1 is recognized at the plasma membrane by *NbPtr1*, the *Nicotiana benthamiana* homologue of *Pseudomonas tomato race 1*. Molecular Plant Pathology, 24, 1312–1318.

Kim, H., Ahn, Y.J., Lee, H., Chung, E.-H., Segonzac, C. & Sohn, K.H. (2023) Diversified host target families mediate convergently evolved effector recognition across plant species. Current Opinion in Plant Biology, 74, 102398.

Kurotani, K., Hirakawa, H., Shirasawa, K., Tagiri, K., Mori, M., Ramadan, A., et al. (2025) Establishing a comprehensive web-based analysis platform for *Nicotiana benthamiana* genome and transcriptome. The Plant Journal, 121, e17178.

Laflamme, B., Dillon, M.M., Martel, A., Almeida, R.N.D., Desveaux, D. & Guttman, D.S. (2020) The pan-genome effector-triggered immunity landscape of a host-pathogen interaction. Science, 367, 763– 768.

Landry, D., González-Fuente, M., Deslandes, L. & Peeters, N. (2020) The large, diverse, and robust arsenal of *Ralstonia solanacearum* type III effectors and their *in planta* functions. Molecular Plant Pathology, 21, 1377–1388.

Lauber, E., González-Fuente, M., Escouboué, M., Vicédo, C., Luneau, J.S., Pouzet, C., et al. (2024) Bacterial host adaptation through sequence and structural variations of a single type III effector gene. iScience, 27, 109224.

Le Roux, C., Huet, G., Jauneau, A., Camborde, L., Trémousaygue, D., Kraut, A., et al. (2015) A receptor pair with an integrated decoy converts pathogen disabling of transcription factors to immunity. Cell, 161, 1074–1088.

Lee, A.H.-Y., Hurley, B., Felsensteiner, C., Yea, C., Ckurshumova, W., Bartetzko, V., et al. (2012) A bacterial acetyltransferase destroys plant microtubule networks and blocks secretion. PLOS Pathogens, 8, e1002523.

Lewis, J.D., Lee, A., Ma, W., Zhou, H., Guttman, D.S. & Desveaux, D. (2011) The YopJ superfamily in plant-associated bacteria. Molecular Plant Pathology, 12, 928–937.

Lewis, J.D., Lee, A.H.-Y., Hassan, J.A., Wan, J., Hurley, B., Jhingree, J.R., et al. (2013) The *Arabidopsis* ZED1 pseudokinase is required for ZAR1-mediated immunity induced by the *Pseudomonas syringae* type III effector HopZ1a. Proceedings of the National Academy of Sciences, 110, 18722–18727.

Lewis, J.D., Wu, R., Guttman, D.S. & Desveaux, D. (2010) Allele-specific virulence attenuation of the *Pseudomonas syringae* HopZ1a type III effector via the *Arabidopsis* ZAR1 resistance protein. PLOS Genetics, 6, e1000894.

Ma, K.-W., Jiang, S., Hawara, E., Lee, D., Pan, S., Coaker, G., et al. (2015) Two serine residues in *Pseudomonas syringae* effector HopZ1a are required for acetyltransferase activity and association with the host co-factor. New Phytologist, 208, 1157–1168.

Ma, K.-W. & Ma, W. (2016) YopJ family effectors promote bacterial infection through a unique acetyltransferase activity. Microbiology and Molecular Biology Reviews, 80, 1011–1027.

Ma, W., Dong, F.F.T., Stavrinides, J. & Guttman, D.S. (2006) Type III effector diversification via both pathoadaptation and horizontal transfer in response to a coevolutionary arms race. PLOS Genetics, 2, e209.

Martel, A., Laflamme, B., Seto, D., Bastedo, D.P., Dillon, M.M., Almeida, R.N.D., et al. (2020) Immunodiversity of the *Arabidopsis* ZAR1 NLR is conveyed by receptor-like cytoplasmic kinase sensors. Frontiers in Plant Science, 11, 1290.

Mazo-Molina, C., Mainiero, S., Haefner, B.J., Bednarek, R., Zhang, J., Feder, A., et al. (2020) Ptr1 evolved convergently with RPS2 and Mr5 to mediate recognition of AvrRpt2 in diverse *solanaceous* species. The Plant Journal, 103, 1433–1445.

Ngou, B.P.M., Ding, P. & Jones, J.D.G. (2022) Thirty years of resistance: Zig-zag through the plant immune system. The Plant Cell, 34, 1447–1478.

Orth, K., Xu, Z., Mudgett, M.B., Bao, Z.Q., Palmer, L.E., Bliska, J.B., et al. (2000) Disruption of signaling by *Yersinia* Effector YopJ, a ubiquitin-like protein protease. Science, 290, 1594–1597.

Pandey, A., Moon, H., Choi, S., Yoon, H., Prokchorchik, M., Jayaraman, J., et al. (2021) *Ralstonia solanacearum* type III effector RipJ triggers bacterial wilt resistance in *Solanum pimpinellifolium*. Molecular Plant Microbe Interactions, 34, 962–972.

Peeters, N., Carrère, S., Anisimova, M., Plener, L., Cazalé, A.-C. & Genin, S. (2013) Repertoire, unified nomenclature and evolution of the Type III effector gene set in the *Ralstonia solanacearum* species complex. BMC Genomics, 14, 859.

Rufián, J.S., Rueda-Blanco, J., López-Márquez, D., Macho, A.P., Beuzón, C.R. & Ruiz-Albert, J. (2021) The bacterial effector HopZ1a acetylates MKK7 to suppress plant immunity. New Phytologist, 231, 1138–1156.

Sabbagh, C.R.R., Carrere, S., Lonjon, F., Vailleau, F., Macho, A.P., Genin, S., et al. (2019) Pangenomic type III effector database of the plant pathogenic *Ralstonia spp*. PeerJ, 7, e7346.

Sarris, P.F., Duxbury, Z., Huh, S.U., Ma, Y., Segonzac, C., Sklenar, J., et al. (2015) A plant immune receptor detects pathogen effectors that target WRKY transcription factors. Cell, 161, 1089–1100.

Savary, S., Willocquet, L., Pethybridge, S.J., Esker, P., McRoberts, N. & Nelson, A. (2019) The global burden of pathogens and pests on major food crops. Nature Ecology & Evolution, 3, 430–439.

Schultink, A., Qi, T., Bally, J. & Staskawicz, B. (2019) Using forward genetics in *Nicotiana benthamiana* to uncover the immune signaling pathway mediating recognition of the *Xanthomonas perforans* effector XopJ4. New Phytologist, 221, 1001–1009.

Seto, D., Koulena, N., Lo, T., Menna, A., Guttman, D.S. & Desveaux, D. (2017) Expanded type III effector recognition by the ZAR1 NLR protein using ZED1-related kinases. Nature Plants, 3, 1–4.

Szczesny, R., Büttner, D., Escolar, L., Schulze, S., Seiferth, A. & Bonas, U. (2010) Suppression of the AvrBs1-specific hypersensitive response by the YopJ effector homolog AvrBsT from *Xanthomonas* depends on a SNF1-related kinase. New Phytologist, 187, 1058–1074.

Tasset, C., Bernoux, M., Jauneau, A., Pouzet, C., Brière, C., Kieffer-Jacquinod, S., et al. (2010) Autoacetylation of the *Ralstonia solanacearum* effector PopP2 targets a lysine residue essential for RRS1-R-mediated immunity in *Arabidopsis*. PLOS Pathogens, 6, e1001202.

Teper, D., Girija, A.M., Bosis, E., Popov, G., Savidor, A. & Sessa, G. (2018) The *Xanthomonas euvesicatoria* type III effector XopAU is an active protein kinase that manipulates plant MAP kinase signaling. PLOS Pathogens, 14, e1006880.

Thieme, F., Szczesny, R., Urban, A., Kirchner, O., Hause, G. & Bonas, U. (2007) New type III effectors from *Xanthomonas campestris pv. vesicatoria* trigger plant reactions dependent on a conserved N-myristoylation motif. Molecular Plant Microbe Interactions, 20, 1250–1261.

Traore, S.M., Eckshtain-Levi, N., Miao, J., Castro Sparks, A., Wang, Z., Wang, K., et al. (2019) *Nicotiana* species as surrogate host for studying the pathogenicity of *Acidovorax citrulli*, the causal agent of bacterial fruit blotch of cucurbits. Molecular Plant Pathology, 20, 800–814.

Üstün, S., Bartetzko, V. & Börnke, F. (2013) The *Xanthomonas campestris* type III effector XopJ targets the host cell proteasome to suppress salicylic-acid mediated plant defence. PLoS Pathogens, 9, e1003427.

Üstün, S. & Börnke, F. (2015) The *Xanthomonas campestris* type III effector XopJ proteolytically degrades proteasome subunit RPT6. Plant Physiology, 168, 107–119.

Üstün, S., König, P., Guttman, D.S. & Börnke, F. (2014) HopZ4 from *Pseudomonas syringae*, a member of the HopZ type III effector family from the YopJ superfamily, inhibits the proteasome in plants. Molecular Plant Microbe Interactions, 27, 611–623.

Vinatzer, B.A., Teitzel, G.M., Lee, M.-W., Jelenska, J., Hotton, S., Fairfax, K., et al. (2006) The type III effector repertoire of *Pseudomonas syringae pv. syringae* B728a and its role in survival and disease on host and non-host plants. Molecular Microbiology, 62, 26–44.

Wang, Jizong, Hu, M., Wang, Jia, Qi, J., Han, Z., Wang, G., et al. (2019) Reconstitution and structure of a plant NLR resistosome conferring immunity. Science, 364, eaav5870.

Xia, Y., Zou, R., Escouboué, M., Zhong, L., Zhu, C., Pouzet, C., et al. (2021) Secondary-structure switch regulates the substrate binding of a YopJ family acetyltransferase. Nature Communications, 12, 5969.

Zhang, Z.-M., Ma, K.-W., Yuan, S., Luo, Y., Jiang, S., Hawara, E., et al. (2016) Structure of a pathogen effector reveals the enzymatic mechanism of a novel acetyltransferase family. Nature Structural & Molecular Biology, 23, 847–852.

Zhou, H., Morgan, R., Guttman, D. & Ma, W. (2009) Allelic variants of the *Pseudomonas syringae* type III effector HopZ1 are differentially recognized by plant resistance systems. Molecular Plant Microbe Interactions, 22, 176–89.

