## Supplementary material for "The recognition of YopJ family effectors depends on ZAR1/JIM2 immune complex in *Nicotiana benthamiana*": Kim I et al_supp info

### Figure S1

| Bacterial genus | Species/strain | Effector | ID number | HR in *N.benthamiana*[1)] |
|---|---|---|---|---|
| *Pseudomonas* | *syringae pv. syringae* | HopZ1a | AAR02168 | O, Ma et al., 2006 |
| | *syringae pv. savastanoi* | HopZ1b | WP_004661226 | O, Ma et al., 2006 |
| | *syringae pv. maculicola* | HopZ1c | AAL84244 | X, Ma et al., 2006 |
| | *syringae pv. syringae* | HopZ2 | ABK13722.1 | O, Ma et al., 2006 |
| | *syringae pv. syringae* | HopZ3 | AAF71492 | X, Ma et al., 2006 |
| | *syringae pv. lachrymans* | HopZ4 | WIO57394.1 | O, this study |
| | *syringae pv. actinidiae* | HopZ5 | AKT29515.1 | O, Ma et al., 2006 |
| *Xanthomonas* | *euvesicatoria* | XopJ1 | CAJ23833 | O, Thieme et al., 2007 |
| | *euvesicatoria* | XopJ2 (AvrBsT) | WP_074052319.1 | O, Orth et al., 2000 |
| | *euvesicatoria pv. vesicatoria str. 85-10* | XopJ3 (AvrRxv) | CAJ22102 | O, Albers et al., 2019 |
| | *perforans* | XopJ4 (AvrXv4) | WP_008572727.1 | O, Schultink et al., 2019 |
| | *compestris pv. campestris* | XopJ5 (AvrXccB) | AAM42989.1 | X, this study |
| | *compestris pv. campestris* | XopJ6 | ND | X, this study |
| *Ralstonia* | *pseudosolanacearum Pe_9* | RipJ | CAD15839 | X, this study |
| | *solanacearum CFBP2957* | RipK | RCFBP_mp10024 | X, this study |
| | *solanacearum* | RipP1 (PopP1) | CAF32331 | O, Poueymiro et al., 2009 |
| | *pseudosolanacearum GMI 1000* | RipP2 (PopP2) | CAD14570 | X, Sohn et al, 2014 |
| *Acidovorax* | *citrulli AAC00-1* | Aave2166 | ABM32744 | O, Traore et al., 2019 |
| | *citrulli AAC00-1* | Aave2708 | ABM33278 | O, Traore et al., 2019 |
| *Erwinia* | *amylovora* | Eop1 | AAF63400 | X, this study |

[1)] Cell death occurrence (O) or absence (X)

### Figure S2

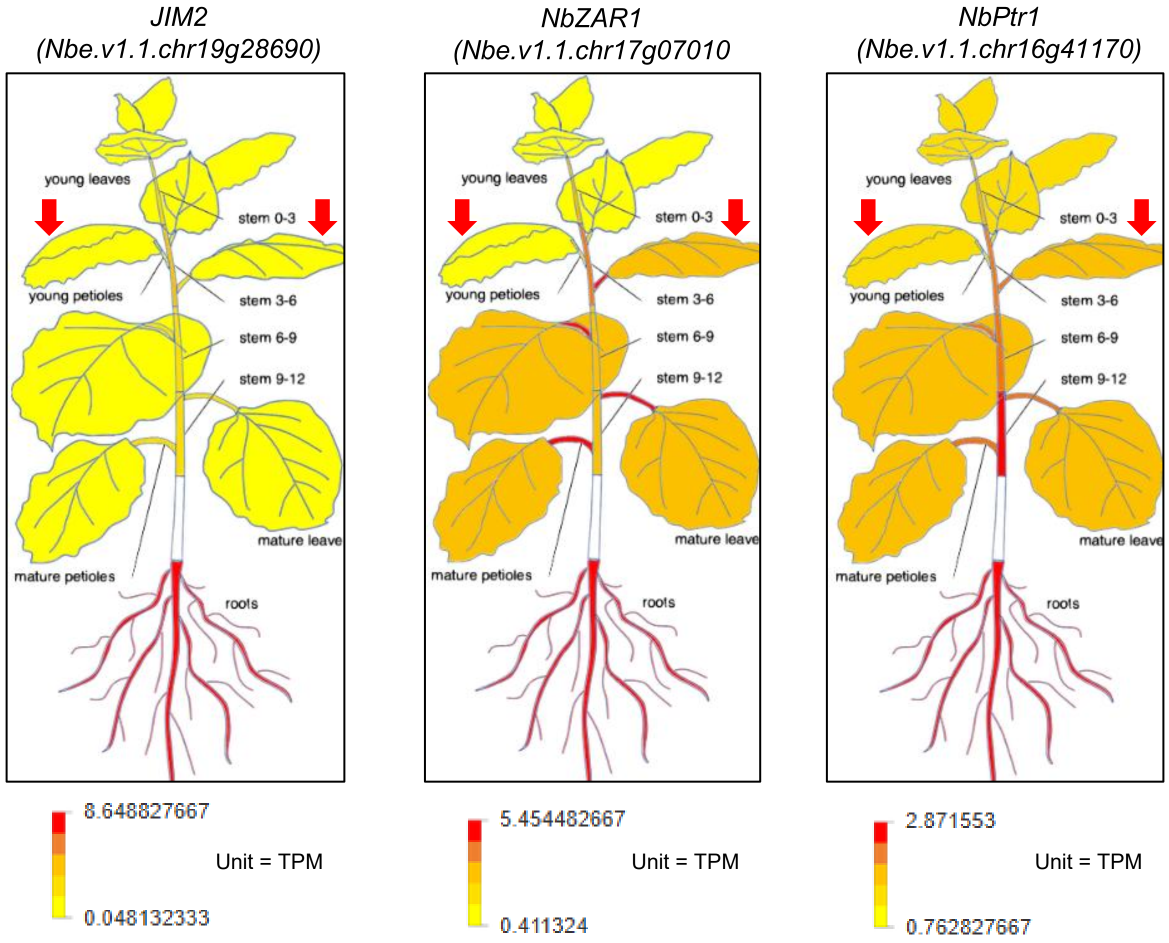
JIM2
(Nbe.v1.1.chr19g28690)
young leaves
stem 0-3
young petioles
stem 3-6
stem 6-9
stem 9-12
mature leave
mature petioles
roots
8.648827667
Unit = TPM
0.048132333
NbZAR1
(Nbe.v1.1.chr17g07010
young leaves
stem 0-3
young petioles
stem 3-6
stem 6-9
stem 9-12
mature leave
mature petioles
roots
5.454482667
Unit = TPM
0.411324
NbPtr1
(Nbe.v1.1.chr16g41170)
young leaves
stem 0-3
young petioles
stem 3-6
stem 6-9
stem 9-12
mature leave
mature petioles
roots
2.871553
Unit = TPM
0.762827667

### Figure S3

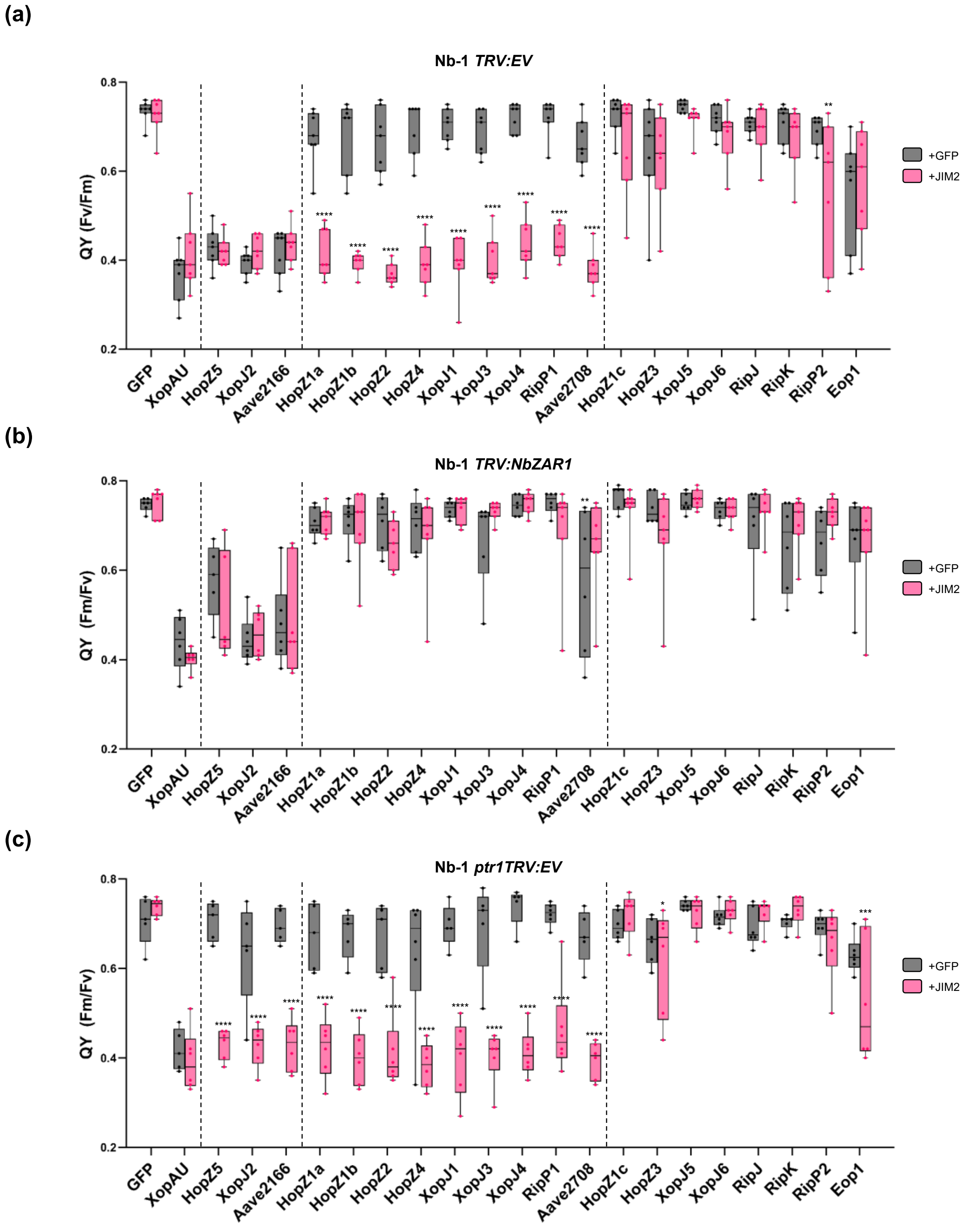
(a)
Nb-1 TRV:EV
QY (Fv/Fm)
+GFP
+JIM2
GFP
XopAU
HopZ5
XopJ2
Aave2166
HopZ1a
HopZ1b
HopZ2
HopZ4
XopJ1
XopJ3
XopJ4
RipP1
Aave2708
HopZ1c
HopZ3
XopJ5
XopJ6
RipJ
RipK
RipP2
Eop1
(b)
Nb-1 TRV:NbZAR1
QY (Fm/Fv)
(c)
Nb-1 ptr1TRV:EV
QY (Fm/Fv)

### Figure S4

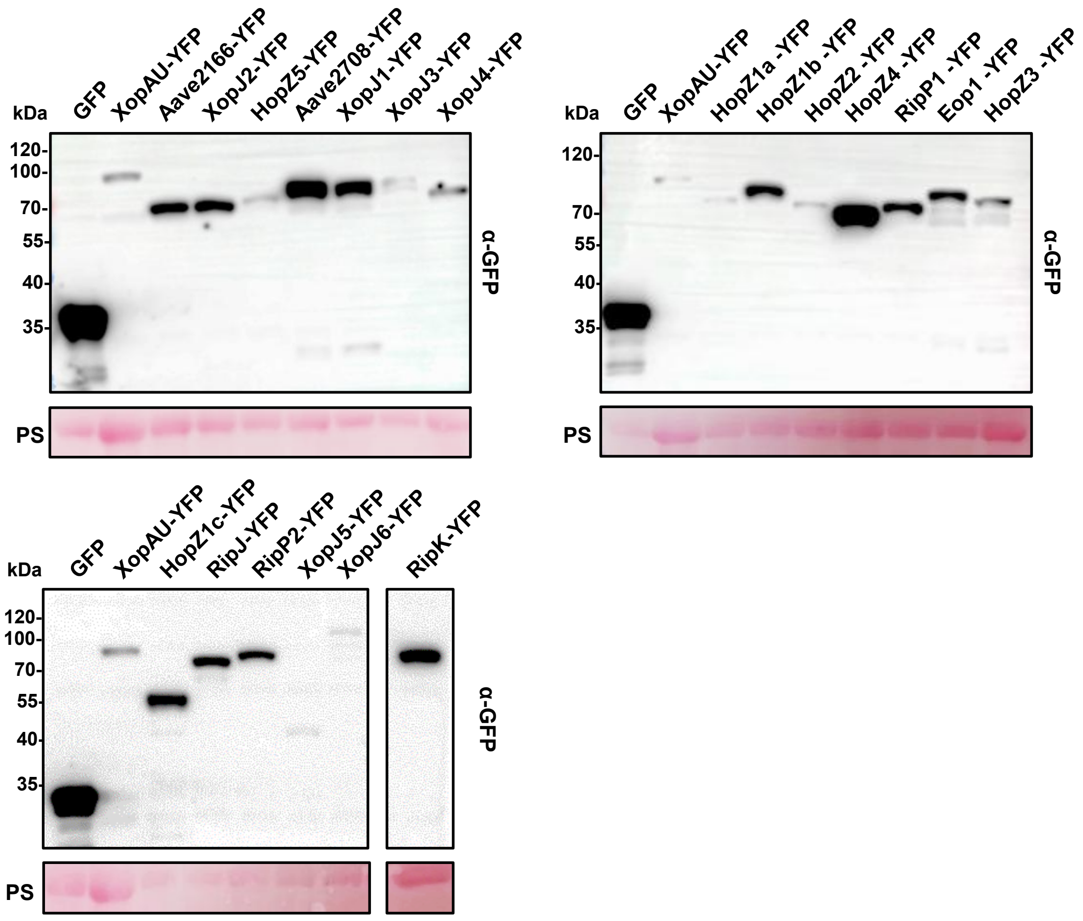
kDa
GFP
XopAU-YFP
Aave2166-YFP
XopJ2-YFP
HopZ5-YFP
Aave2708-YFP
XopJ1-YFP
XopJ3-YFP
XopJ4-YFP
120-
100-
70-
55-
40-
35-
α-GFP
PS
kDa
GFP
XopAU-YFP
HopZ1a -YFP
HopZ1b -YFP
HopZ2 -YFP
HopZ4 -YFP
RipP1 -YFP
Eop1 -YFP
HopZ3 -YFP
120-
70-
55-
40-
35-
α-GFP
PS
kDa
GFP
XopAU-YFP
HopZ1c-YFP
RipJ-YFP
RipP2-YFP
XopJ5-YFP
XopJ6-YFP
RipK-YFP
120-
100-
70-
55-
40-
35-
α-GFP
PS

### Figure S5

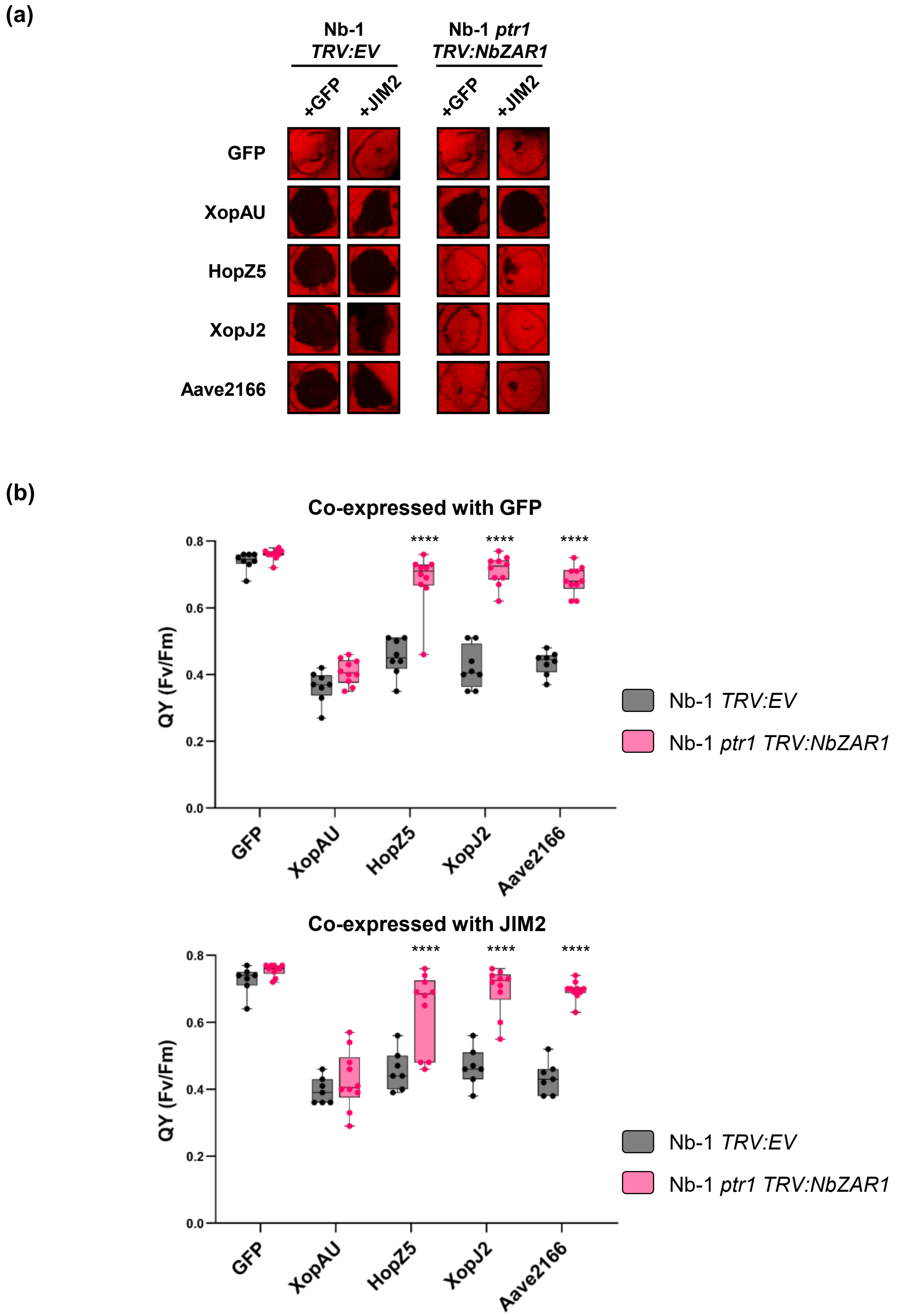

(a)
Nb-1
TRV:EV
Nb-1 ptr1
TRV:NbZAR1
+GFP
+JIM2
+GFP
+JIM2
GFP
XopAU
HopZ5
XopJ2
Aave2166
(b)
Co-expressed with GFP
QY (Fv/Fm)
0.0
0.2
0.4
0.6
0.8
****
****
****
GFP
XopAU
HopZ5
XopJ2
Aave2166
Nb-1 TRV:EV
Nb-1 ptr1 TRV:NbZAR1
Co-expressed with JIM2
QY (Fv/Fm)
0.0
0.2
0.4
0.6
0.8
****
****
****
GFP
XopAU
HopZ5
XopJ2
Aave2166
Nb-1 TRV:EV
Nb-1 ptr1 TRV:NbZAR1

### Figure S6

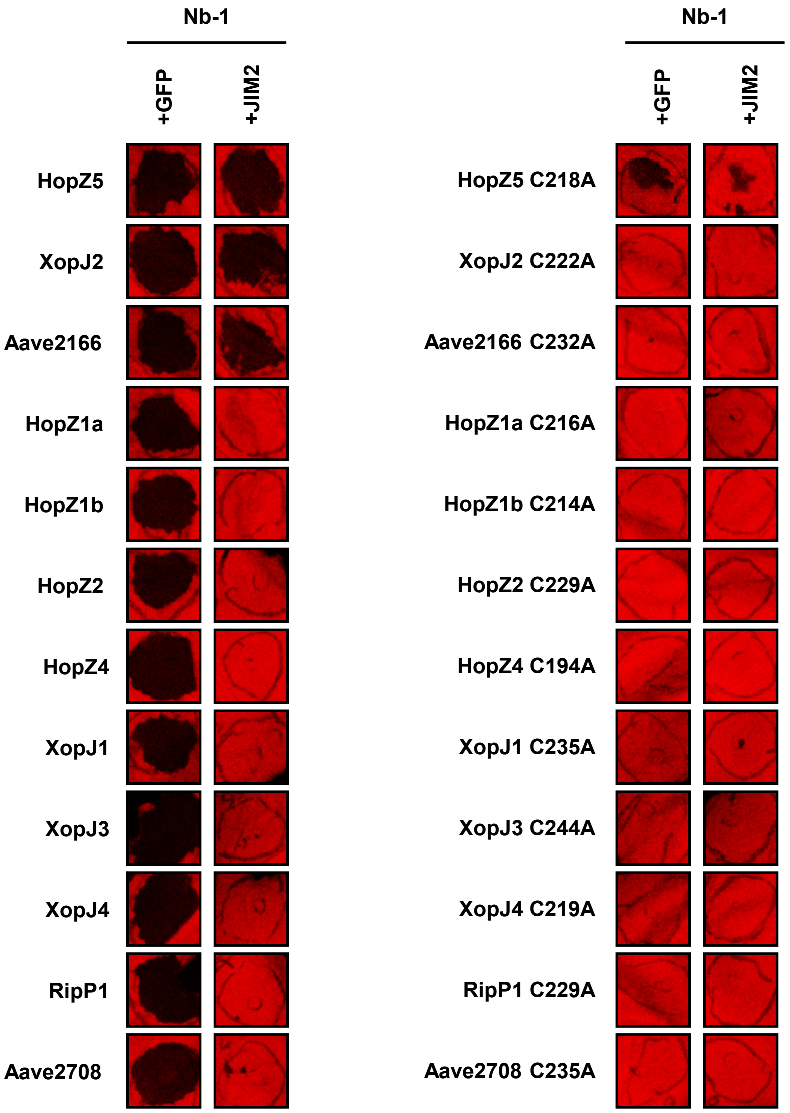

Nb-1
+GFP
+JIM2
HopZ5
XopJ2
Aave2166
HopZ1a
HopZ1b
HopZ2
HopZ4
XopJ1
XopJ3
XopJ4
RipP1
Aave2708
Nb-1
+GFP
+JIM2
HopZ5 C218A
XopJ2 C222A
Aave2166 C232A
HopZ1a C216A
HopZ1b C214A
HopZ2 C229A
HopZ4 C194A
XopJ1 C235A
XopJ3 C244A
XopJ4 C219A
RipP1 C229A
Aave2708 C235A

### Figure S7

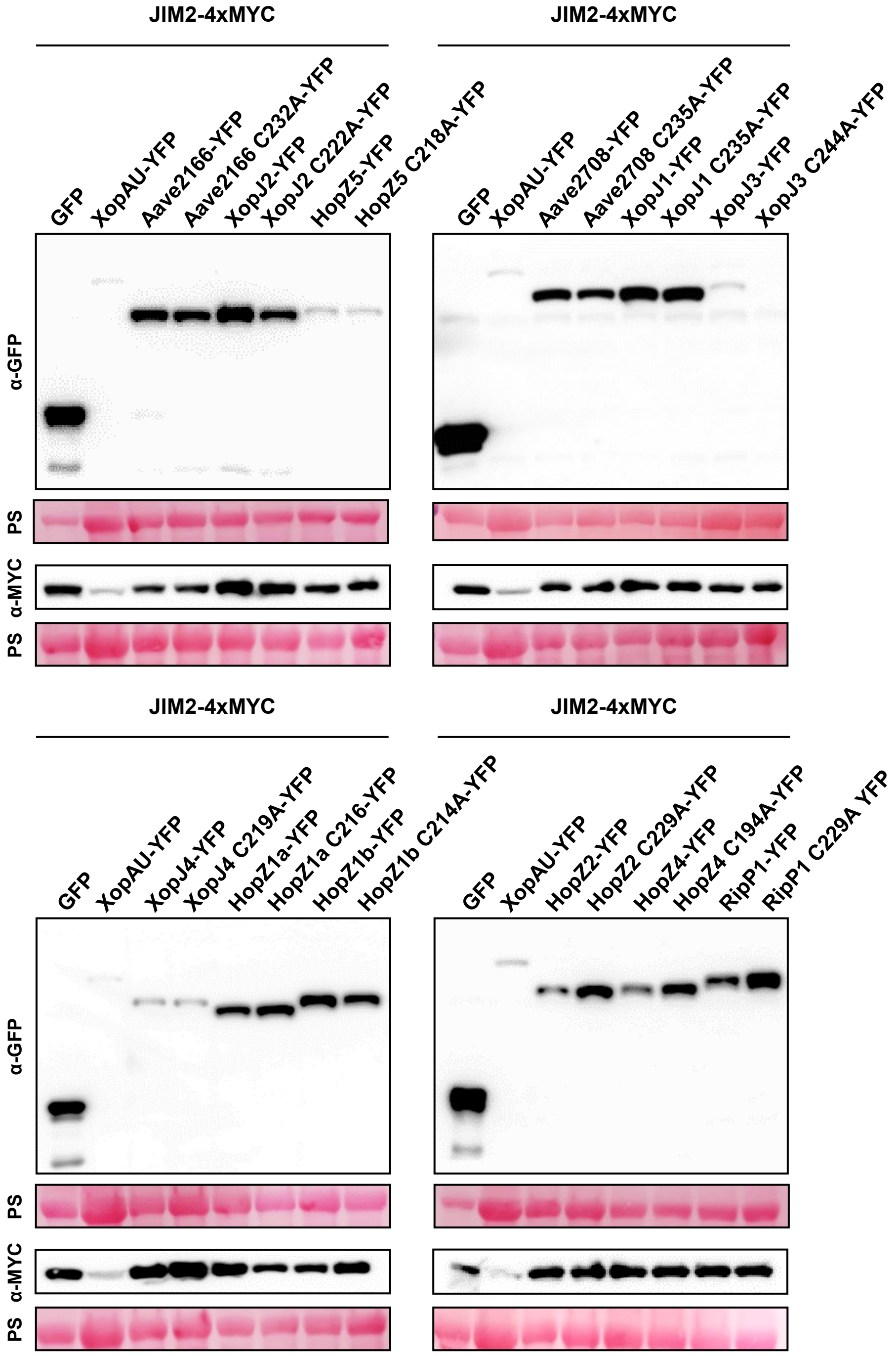
JIM2-4xMYC
GFP
XopAU-YFP
Aave2166-YFP
Aave2166 C232A-YFP
XopJ2-YFP
XopJ2 C222A-YFP
HopZ5-YFP
HopZ5 C218A-YFP
α-GFP
PS
α-MYC
PS
JIM2-4xMYC
GFP
XopAU-YFP
Aave2708-YFP
Aave2708 C235A-YFP
XopJ1-YFP
XopJ1 C235A-YFP
XopJ3-YFP
XopJ3 C244A-YFP
JIM2-4xMYC
GFP
XopAU-YFP
XopJ4-YFP
XopJ4 C219A-YFP
HopZ1a-YFP
HopZ1a C216-YFP
HopZ1b-YFP
HopZ1b C214A-YFP
α-GFP
PS
α-MYC
PS
JIM2-4xMYC
GFP
XopAU-YFP
HopZ2-YFP
HopZ2 C229A-YFP
HopZ4-YFP
HopZ4 C194A-YFP
RipP1-YFP
RipP1 C229A YFP

### Figure S8

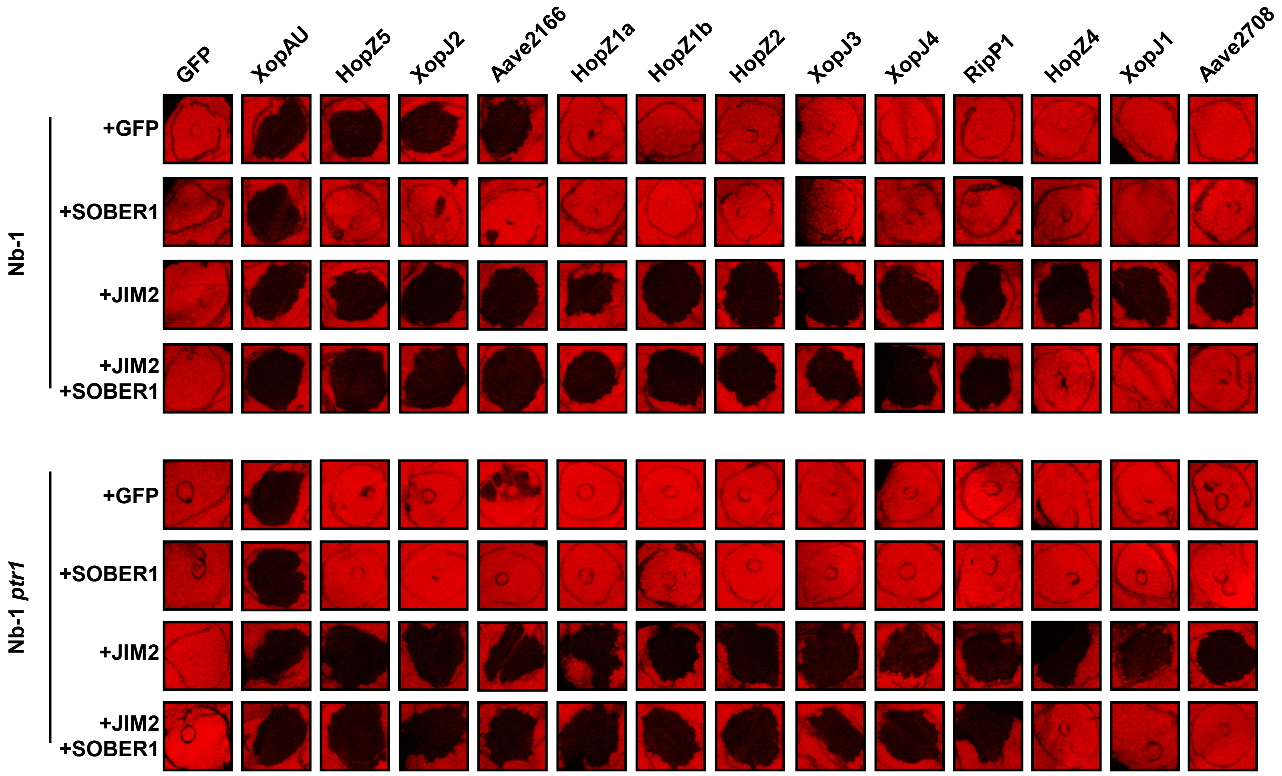
GFP
XopAU
HopZ5
XopJ2
Aave2166
HopZ1a
HopZ1b
HopZ2
XopJ3
XopJ4
RipP1
HopZ4
XopJ1
Aave2708
Nb-1
+GFP
+SOBER1
+JIM2
+JIM2
+SOBER1
Nb-1 *ptr1*
+GFP
+SOBER1
+JIM2
+JIM2
+SOBER1

### Figure S9

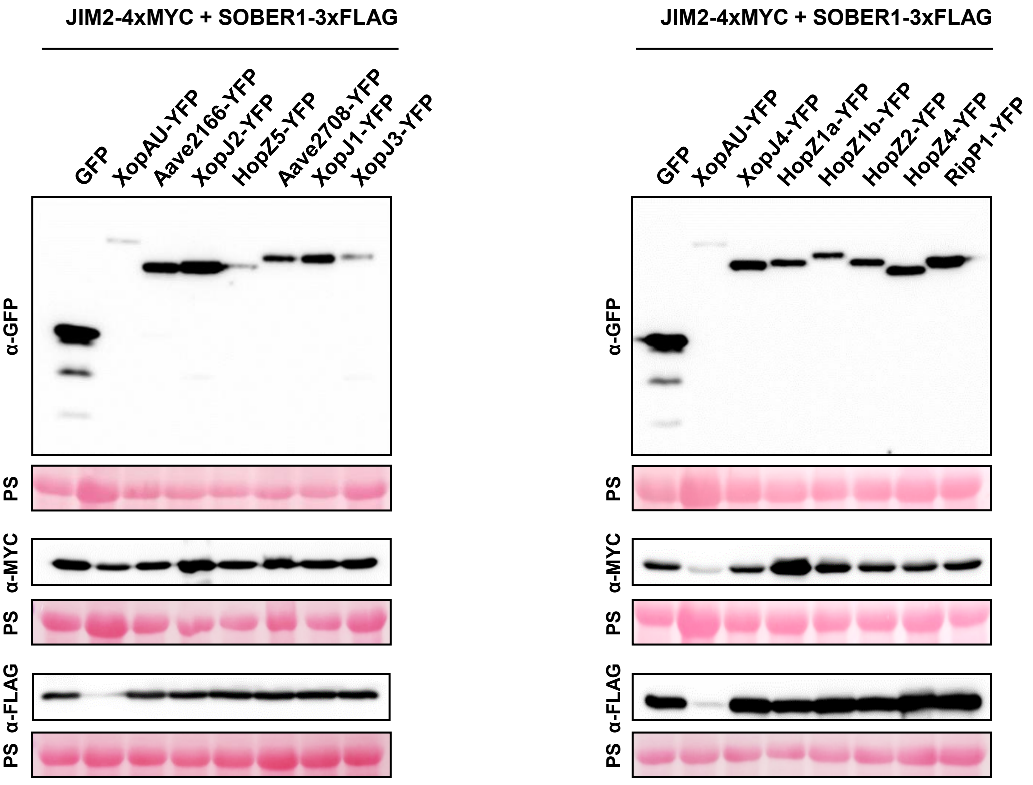
JIM2-4xMYC + SOBER1-3xFLAG
GFP
XopAU-YFP
Aave2166-YFP
XopJ2-YFP
HopZ5-YFP
Aave2708-YFP
XopJ1-YFP
XopJ3-YFP
α-GFP
PS
α-MYC
PS
α-FLAG
PS
JIM2-4xMYC + SOBER1-3xFLAG
GFP
XopAU-YFP
XopJ4-YFP
HopZ1a-YFP
HopZ1b-YFP
HopZ2-YFP
HopZ4-YFP
RipP1-YFP
α-GFP
PS
α-MYC
PS
α-FLAG
PS
